# Mucosal microbiomes and *Fusobacterium* genomics in Vietnamese colorectal cancer patients

**DOI:** 10.1101/2022.02.25.481918

**Authors:** Hoang N. H. Tran, Trang Nguyen Hoang Thu, Phu Huu Nguyen, Chi Nguyen Vo, Khanh Van Doan, Chau Nguyen Ngoc Minh, Ngoc Tuan Nguyen, Van Ngoc Duc Ta, Khuong An Vu, Thanh Danh Hua, To Nguyen Thi Nguyen, Tan Trinh Van, Trung Pham Duc, Ba Lap Duong, Phuc Minh Nguyen, Vinh Chuc Hoang, Duy Thanh Pham, Guy E. Thwaites, Lindsay J. Hall, Daniel J. Slade, Stephen Baker, Vinh Hung Tran, Hao Chung The

## Abstract

Perturbations in the gut microbiome have been linked to the promotion and prognosis of colorectal cancer (CRC), with the colonic overabundance of *Fusobacterium nucleatum* shown as the most consistent marker. Despite the increasing health burden inflicted by CRC in low- and middle-income countries like Vietnam, the CRC-specific microbiome in these populations remains underexplored. Here we conducted a study in Vietnam to enrol 43 CRC patients (cases) and 25 patients with non-cancerous colorectal polyps (controls) between December 2018 and January 2020. Our study investigated the mucosal microbiome signature and genomic diversity of *Fusobacterium* in Vietnamese CRC patients, using a combination of 16S rRNA gene profiling, anaerobic microbiology, and whole genome sequencing. We found that several oral bacteria, including *F. nucleatum* and *Leptotrichia*, were significantly more abundant in the tumour mucosa, and these two bacteria were also more enriched in tumours of advanced CRC stages (III-IV). We obtained 53 *Fusobacterium* genomes from the saliva, tumour and non-tumour mucosa of six CRC patients. Isolates from the gut mucosa belonged to diverse *F. nucleatum* subspecies (*nucleatum, animalis, vincentii, polymorphum*) and a potential new subspecies of *F. periodonticum*. The *Fusobacterium* population within each individual was distinct and in many cases diverse, with minimal intra-clonal variation. Phylogenetic analyses showed that within each individual, tumour-associated *Fusobacterium* were clonal to those isolated from non-tumour mucosa, but distantly related to those isolated from saliva. Genes encoding major virulence factors (Fap2 and RadD) showed variability in length and evidence of horizontal gene transfer. Our work provides a framework to understand the genomic diversity of *Fusobacterium* within the CRC patients, which can be exploited for the development of CRC diagnostic and therapeutic options targeting this oncobacterium.

## Introduction

Colorectal cancer (CRC) is the second leading cause of cancer mortality worldwide, contributing to an estimate of 850,000 deaths and ∼1.8 million new cases in 2018 [1,2]. The majority of CRC cases are sporadic (without clear heredity components), with well-established lifestyle risk factors attributed to obesity, alcohol consumption and a diet enriched with red or processed meat [3]. The vast and diverse microbial community inhabiting the colon (termed the gut microbiome) is an integral part of human health, and act as an important interface mediating the interactions between environmental cues, host biology, and CRC [4,5]. Research on CRC gut microbiome has consistently underlined the abundances of certain marker bacteria, among which *Fusobacterium nucleatum* has been most widely reported and intensively studied [6–10].

The Gram-negative rod-shaped *F. nucleatum* is a common anaerobic member of the human oral microbiome, and it is currently composed of four subspecies (*nucleatum, vincentii, animalis*, and *polymorphum*) [11]. Mechanistic studies have demonstrated that *F. nucleatum* possesses several virulence factors, most notably FadA and Fap2, which enable the bacteria to potentiate colonic tumourigenesis. The adhesin FadA binds to E-cadherin in CRC cells and activates the β-catenin-dependent oncogenic pathways [12], while the lectin Fap2 further facilitates *F. nucleatum* invasion into CRC cells by specifically binding to the tumour-enriched carbohydrate Gal-GalNAc [13]. Such interaction triggers the secretions of the pro-inflammatory (IL-8) and pro-metastatic (CXCL-1) cytokines, creating a tumour environment conditioned for accelerated growth and migratory tendency [14]. Recent studies have further highlighted that the bacteria could induce DNA damage in oral and colorectal cancerous cells [15,16]. Additionally, *F. nucleatum* lipopolysaccharide was shown to induce resistance to chemotherapy via activation of the autophagy machinery in CRC cells, thus complicating effective CRC treatment [17]. As a result, enrichment of *F. nucleatum* in CRC microbiomes has been associated with more severe prognosis and poorer overall survival, particularly in a subset of patients with mesenchymal tumours [18– 20]. Preclinical research demonstrated that *F. nucleatum* elimination by antibiotics reduced colorectal tumour proliferation in mice [21]. These evidences strongly support for the utilization of *F. nucleatum* as a target for CRC diagnosis and therapy, but current translational potential is hampered by the lack of insights into *F. nucleatum* diversity and its genomic characteristics in CRC patients.

The majority of microbiome studies, on either healthy or CRC cohorts, were conducted in high-income countries, and such data are sparse regarding populations in developing settings, where host factors, diet and lifestyle could greatly influence the gut microbiome composition and function. Vietnam has an increasing ageing population adopting a more ‘Westernized’ diet and sedentary lifestyle [22], where CRC incidence is predicted to climb and rank as among the top three cancers by 2025 [23]. Therefore, CRC microbiome studies in Vietnam are necessary to establish the basis for the implementation of microbiome-oriented strategies for CRC prevention, diagnosis, prognosis and therapy. We set out to investigate the microbiome signatures of Vietnamese CRC patients, by applying 16S-rRNA gene profiling on the saliva and gut mucosa collected from patients with CRC and non-cancerous colorectal polyps. Additionally, different from prior studies, we used anaerobic culturing and whole genome sequencing (WGS) to study the genomic diversity of *Fusobacterium* colonizing these CRC patients, allowing an in-depth and high-resolution examination of these bacterial populations.

## Results

### Gut mucosal, but not salivary, microbiomes differ significantly between CRC and controls

We enrolled 43 CRC patients (cases) and 25 patients with colorectal polyps (controls) between December 2018 and January 2020. 16S rRNA microbiome profiling was performed for all the saliva and gut mucosa samples collected from the participants, including tissues originating from the diseased (CRC tumour or polyps) and the adjacent normal sites. To limit the scope of this study, we selected participants with tumours/polyps detected in the distal colon or rectum. The patients’ demographic and clinical data were summarized in Table 1, which showed that there were no significant differences between the two groups. All polyps showed not more than low-grade dysplasia (i.e. non-cancerous), demonstrating the validity of our control group. Microbiome profiling identified 865 filtered amplicon sequence variants (ASVs – a marker for distinct taxonomic classification) among 66 saliva samples, with a median library size of 36,250 paired-end reads [IQR: 31,827 – 50,317]. Due to their lower microbial biomass, the library size of gut mucosal microbiomes was smaller (median: 17,711 [IQR: 9,037 – 30,135]), with 1,073 filtered ASVs detected across 129 mucosa samples (seven removed).

**Table 1.** Baseline characteristics of patients recruited in this study. Overweight/obesity was classified using WHO recommendation for Asian populations. Oral diseases include self-reported gingivitis, periodontitis or halitosis. The number in each cell refers to median with interquartile range in brackets, percentage or count number for each category.

|  | CRC cases (n=42) | Controls (n=21) | p-value |
| --- | --- | --- | --- |
| **Age** | 64 [54 - 69] | 60 [53 - 66] | 0.359 |
| **Male sex** | 62% | 76% | 0.395 |
| **BMI** | 22.9 [20.85 - 24.95] | 22.2 [21.1 - 23.4] | 0.387 |
| **Overweight/obesity** | 47.60% | 33% | 0.409 |
| **Diabetes** | 19% | 19% | 1 |
| **High blood pressure** | 52% | 47.60% | 0.79 |
| **Active smoking in the last two years** | 21.40% | 19% | 1 |
| **Oral diseases** | 33% | 38% | 0.782 |
| **Family history of cancer** | 19% | 19% | 1 |
| **Location of sampled mucosa** |  |  | 0.533 |
| *Descending colon* | 7 | 3 |  |
| *Sigmoid colon* | 28 | 12 |  |
| *Rectum* | 7 | 6 |  |
| **Size of tumour/polyp (cm)** | 5 [4 - 5.75] | 1 [0.7 - 1.2] |  |
| **TNM stage of cancer** | II (18), III (20), IV (4) |  |  |
| **Polyp dysplasia grade** |  | low (4), none (17) |  |

Ordination by principal coordinate analysis (PCoA), based on phylogenetic-assisted isometric log-ratio (PhIRL) transformed value, showed that the salivary microbiomes of CRC and controls completely overlapped (Figure 1A). Only active smoking within the last two years, but not CRC status, was significantly associated with the salivary microbiome structure (RDA, p-value = 0.033). Likewise, only two ASVs belonging to the genera *Leptotrichia* and *Solobacterium* were consistently identified as significantly more abundant in the CRC’s salivary microbiome. These point to the high structural similarity in the salivary microbiome between the two groups. By contrast, the gut mucosal microbiomes differ significantly based on CRC status (Figure 1B). CRC and diabetes significantly contributed to the variance in the gut microbiome (RDA, p-value < 0.05). Gut mucosa collected within a participant (tumour and non-tumour for CRC, biopsy and polyp for control) shared more similarity in their microbiomes than those of the same sample type between participants (Figure 1C), resembling findings from previous research [8]. We also conducted these analyses using the weighted Unifrac and Bray-Curtis distances, which produced similar interpretations. Additionally, we performed unsupervised clustering on gut mucosa microbiomes, which showed the presence of two robust community state types (CSTs) supported by an out-of-bag error rate of 10.8% in a random forest classification. This algorithm also identified that several ‘balances’ (Proteobacteria/Actinobacteria, other bacteria/Lachnospiraceae) contributed significantly in separating the two CSTs (Figure S1). CST1 was generally more enriched in Gammaproteobacteria (mostly *Escherichia*) while CST2 had higher abundance of Actinobacteria (mainly *Collinsella*) and Lachnospiraceae (Figure 1D). The two CSTs were similar in library size (p-value = 0.15, t-test), but different in CRC status (p-value=0.002, Fisher-exact test), with the majority of control samples (72%) belonging to CST1. No other tested covariates were associated with CST grouping. Samples from the same patients mostly shared the same CST membership (90.3%, n=56/62 patients with paired microbiomes), and CRC samples were distributed in both CSTs with different proportion (CST1 = 36, CST2 = 50). These findings suggest that while CRC status mainly explained the dissimilarity observed in the gut mucosal microbiomes, their overall configurations were determined by the dominant presence of Gammaproteobacteria (*Escherichia*), possibly driven by an unknown or stochastic factor.

**Figure 1.**
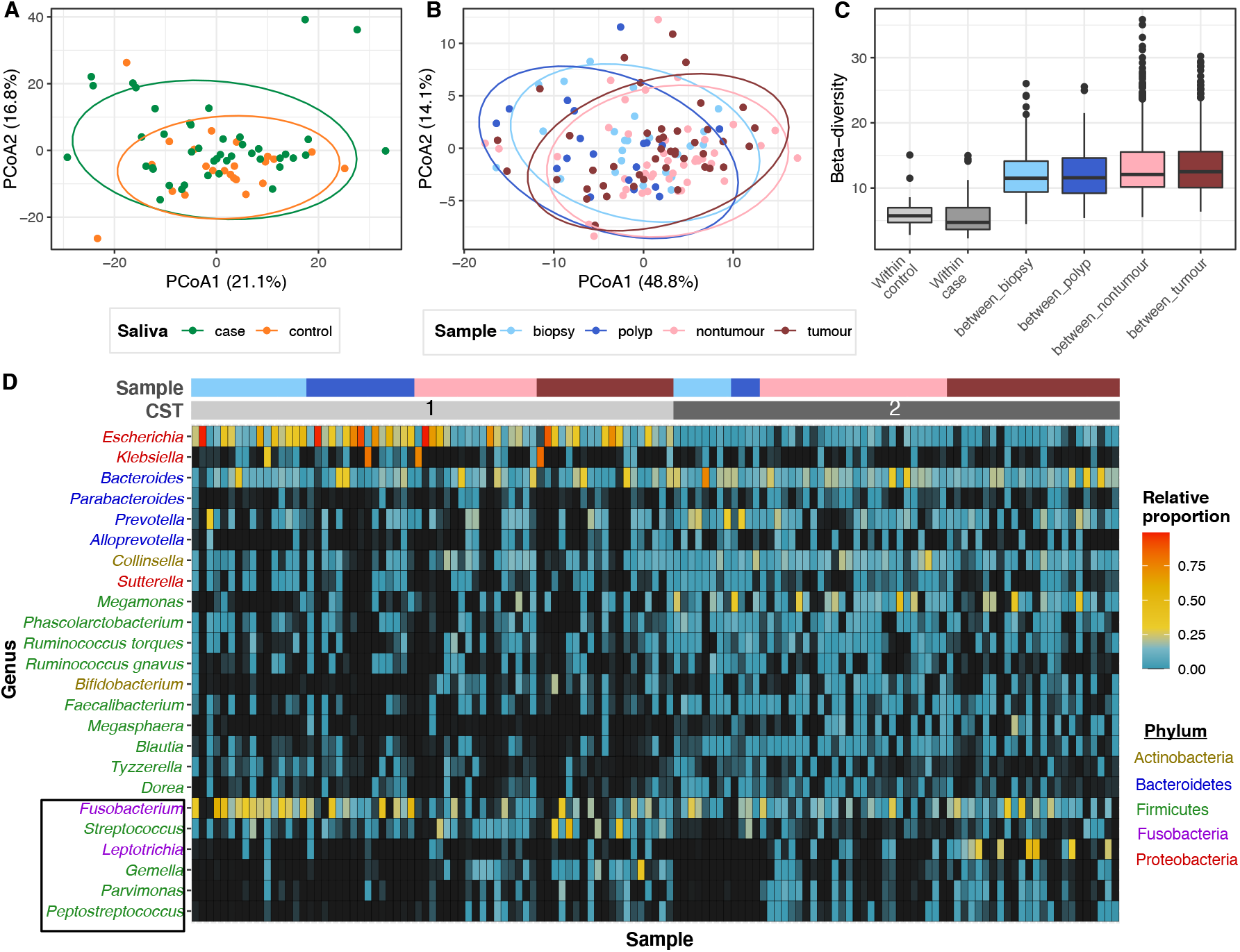
The salivary and gut mucosa microbiomes of colorectal cancer patients. Principal coordinate analyses (PCoA), conducted on phylogenetic-assisted isometric log-ratio (PhILR) transformed data, of (A) 66 salivary microbiomes, and (B) 129 gut mucosa microbiomes, with different CRC groups and sample types denoted by different colours (see Keys; biopsies and polyps collected from controls, nontumours and tumours collected from cases). (C) Boxplot showing the distribution of pairwise beta-diversity, calculated on PhILR transformed values, observed in each gut microbiome category. (D) Heatmap displaying the proportional abundances of 24 most abundant genera, with headers showing the samples’ community state type (CST): CST1 (light gray), CST2 (dark gray), and the corresponding sample type: biopsy (light blue), polyp (dark blue), nontumour (pink), tumour (dark red). Genera were coloured according to their classifications at Phylum level (see Keys). Genera in black box represent ones with probable origin from the oral cavity.

### Enrichment of oral bacteria in the tumour gut mucosa

We applied differential abundance analysis to rigorously detect bacteria enriched in the CRC tumours, by comparing results from different approaches, including ANCOMBC, DESeq2 and corncob (see Methods) [24–26]. Our analyses revealed that ASVs classified as bacteria of putative oral origin (*Gemella, Peptostreptococcus, F. nucleatum, Leptotrichia, Selenomonas sputigena*, and *Campylobacter rectus*) were overabundant in the tumour mucosa, compared to control biopsies (Figure 2B). This finding corroborates the observation that these oral bacteria had a higher relative abundance in CRC gut microbiomes of both CSTs (Figure 1D). Within the CRC patients, tumours also showed an elevated presence of the aforementioned oral bacteria (alongside *Hungatella, Lachnoclostridium*, and *Osillibacter*) when compared to adjacent non-tumour mucosa, albeit with less pronounced fold change (Figure 2A).

**Figure 2.**
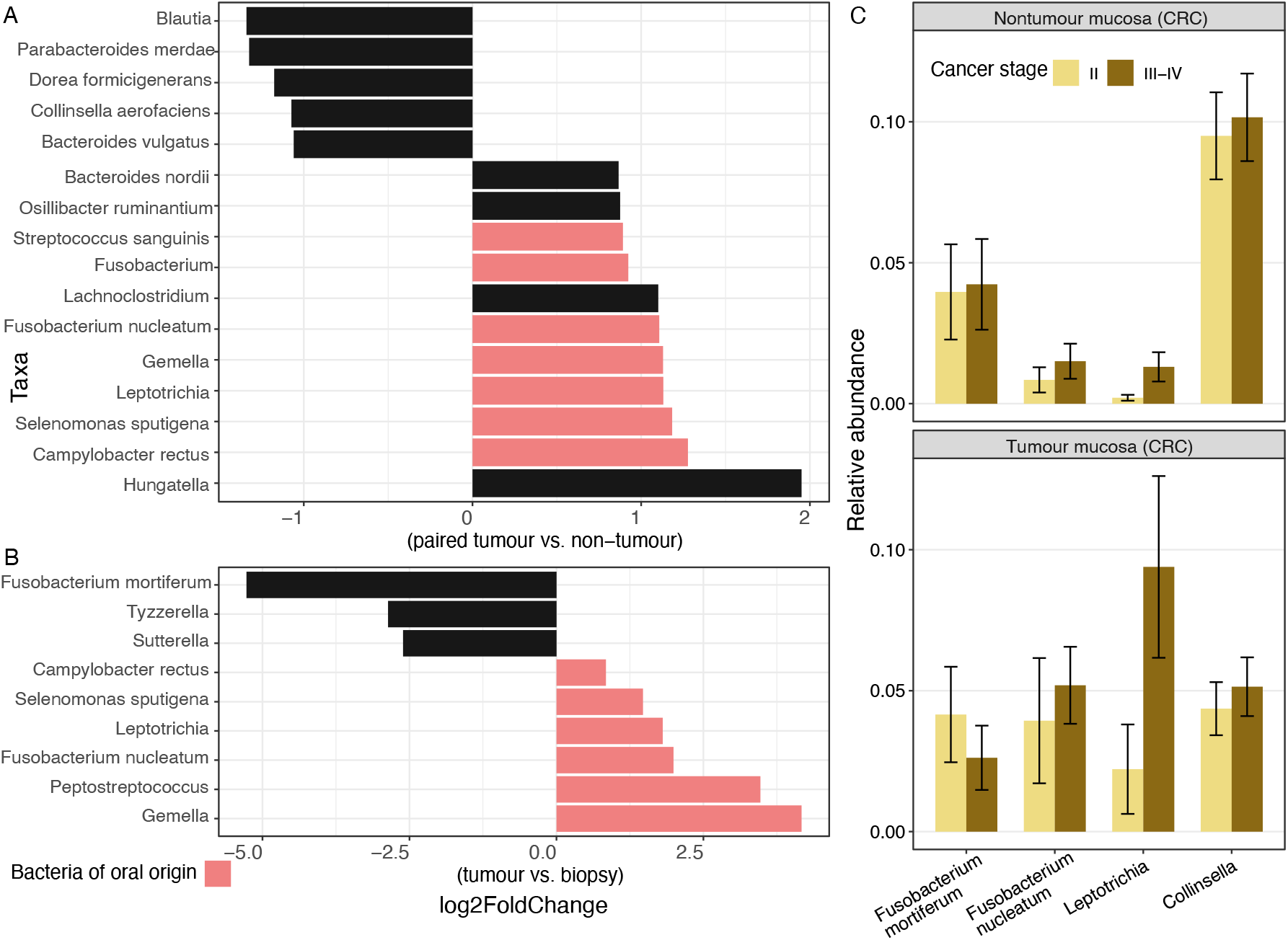
Bacterial taxa significantly abundant among the examined classes. Taxa, or amplicon sequence variants (ASVs), were determined as significant and visualized in (A) and (B) if they were detected in at least two of the three tested approaches (ANCOMBC, DESeq2, corncob; adjusted p-value ≤ 0.05). (A) Log2 fold change of ASVs that differ between paired tumour and non-tumour mucosal microbiomes from case participants, using the full model ‘Patient + sample type’ (n=86). (B) Log2 fold change of ASVs that differ between tumour and biopsy (control) mucosal microbiomes (n=67). Log2 fold change was derived from ANCOMBC test output, and taxa of oral origin were coloured in pink. (C) Relative abundance of ASVs assigned as *Fusobacterium mortiferum* (n=14), *Fusobacterium nucleatum* (n=14), *Leptotrichia* (n=16), and *Collinsella* (n=14) in the tumour and nontumour mucosal microbiomes, stratified by cancer stages (III-IV vs. II).

These increases were coupled with the reduction in abundances of commensal anaerobes in the tumour mucosa, such as *Blautia, Parabacteroides, Dorea*, and *Collinsella*. When comparing between different cancer stages, the increased abundance of one taxon (*Leptotrichia*, ASV-13) was consistently associated with tumours of advanced stages (III-IV), compared to stage II (Figure 2C). Results from DESeq2 alone additionally showed that *F. nucleatum* was also enriched in advanced CRC stages (adjusted p-value <0.05). ASVs confidently assigned as *F. nucleatum* (n=14) and *Leptotrichia* spp. (n=16) were present with at least 0.1% relative abundance in ∼54% and 40% of tumour microbiomes, respectively. On the other hand, tumour samples with low abundances of oral bacteria (<1%) were dominated by other taxa, including *Escherichia/Klebsiella* (n= 4), *Megamonas* (n=4), *Fusobacterium mortiferum* (n=2), and *Helicobacter* (n=1). We performed similar analysis within the control group and showed that only one ASV (*Faecalibacterium*) was consistently depleted in polyps compared to paired biopsies. However, when compared to CRC samples, *F. mortiferum, Tyzzerella*, and *Sutterella* were significantly enriched in the control gut microbiomes (Figure 2B).

To investigate bacterial co-occurrence and their potential interactions, we next used CCLasso and SpiecEasi to construct a correlation network of gut microbiomes from CRC patients (n=86) (Figure 3) [27,28]. Two oral bacteria clusters emerged from this network, one consisting of several *Streptococcus* and *Veillonella* taxa, and another composed mostly of aforementioned tumour-associated ASVs (*Leptotrichia, Selenomonas, F. nucleatum, Streptococcus, Granulicatella, Gemella, Peptostreptococcus*, and *Parvimonas*). The latter cluster exhibited positive correlation with *E. coli*, and antagonism toward *Blautia*, a member of the gut anaerobic commensal network. Besides, other tumour-associated ASVs such as *Hungatella, Lachnoclostridium*, and *C. rectus* were clustered alongside *Negativibacillus* and *Eggerthella*, which showed strong negative correlations with anaerobic gut commensals *Dorea, Bacteroides*, and *Faecalibacterium*. These findings highlight the potential competition between tumour-associated taxa and common gut commensal anaerobes (Lachnospiraceae, Oscillospirales). Other *Fusobacterium* species, *F. mortiferum* and *F. varium* were not linked to the oral clusters, showing that they were mainly gut inhabitants. Comparison with the network constructed from salivary microbiomes revealed that the same tumour-associated ASVs (*F. nucleatum, Gemella, Selenomonas*) formed similar clusters as observed in the CRC gut microbiomes (Figure S2). This indicates that tumour-associated ASVs could have oral origin in this examined cohort, and they likely co-exist in the polymicrobial biofilms (similar to those present in the oral cavity) upon gut colonization. Notably, the tumour-associated *Leptotrichia* (ASV-13) had very low abundance in the salivary microbiome (mean: 0.037%, prevalence: 26%), and it differs from the dominant *Leptotrichia* detected in saliva (ASV-19) in 17 nucleotides (pairwise similarity: 93.3%).

**Figure 3.**
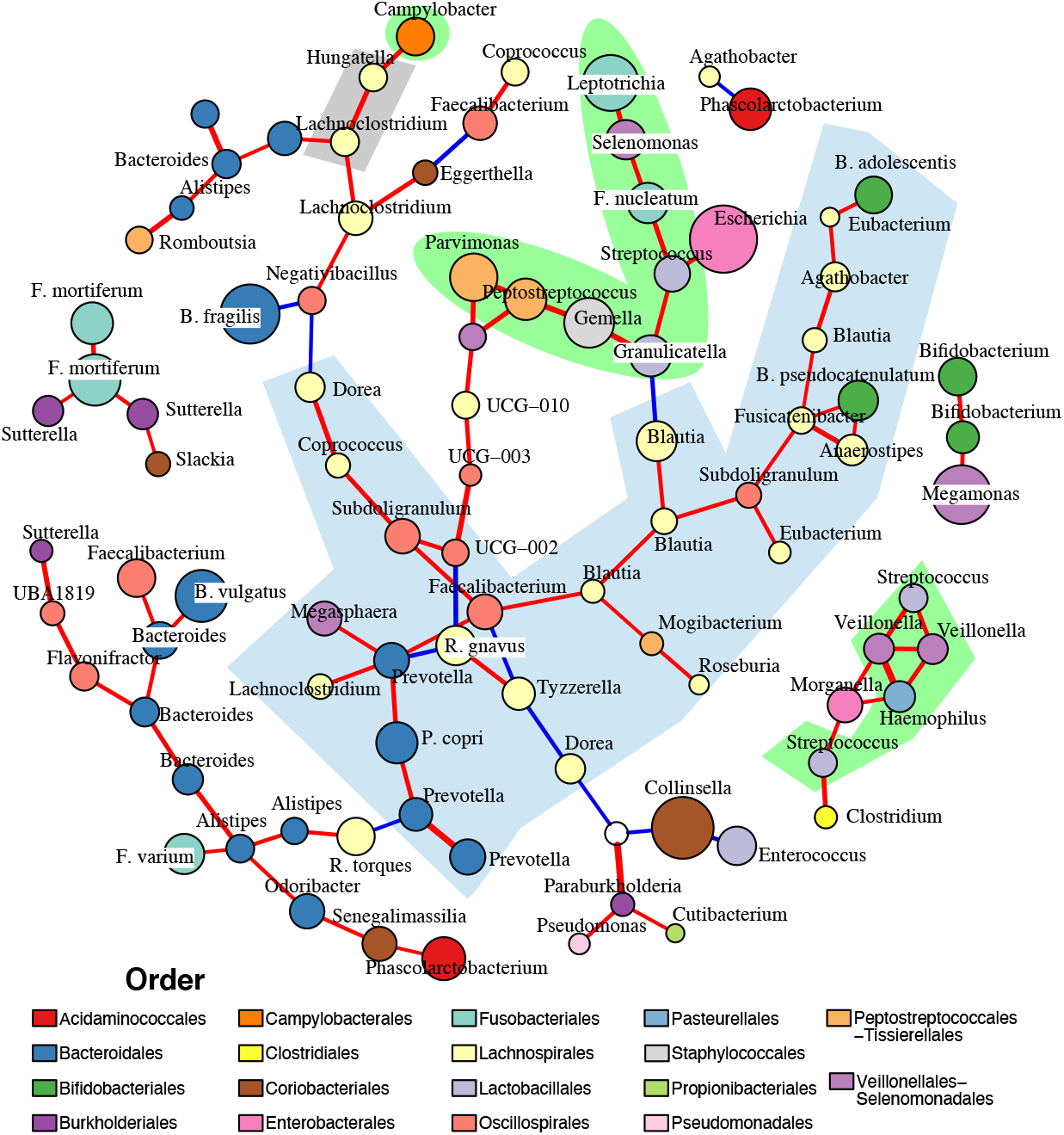
Correlation network of colorectal cancer gut mucosal microbiomes. The network was constructed from 117 most representative ASVs sampled from 86 mucosal microbiomes, outlining significant interactions detected by both CCLasso (p value ≤ 0.01 and absolute correlation strength > 0.37) and SpiecEasi. Positive and negative interactions were coloured as red and blue lines respectively, with line weight proportional to correlation strength. The ASVs (nodes) were coloured based on taxonomic family (see Legend), with sizes proportional to their relative abundances. The light green shaded area entails ASVs identified as members of the human oral microbiome (comparison with expanded Human Oral Microbiome Database); the blue shaded area covers ASVs identified as gut anaerobic commensals (Lachnospirales, Bacteroidales, Bifidobacteriales, Oscillospirales); and gray shaded area covers other tumour-associated ASVs (as identified in Figure 2A).

### Diverse *Fusobacterium* colonizes CRC patients

Since *F. nucleatum* was more enriched in the tumour microbiomes and previously demonstrated to promote tumourigenesis, we next studied the population structure of *Fusobacterium* recovered from CRC patients. Six patients with a *Fusobacterium* relative abundance at the tumour site exceeding 10% (except for patient 18) and covered different cancer stages were selected for *Fusobacterium* isolation. In total, we isolated 56 presumptive *Fusobacterium* organisms, as identified by the matrix-assisted laser desorption/ionization time of flight mass spectrometer (MALDI-TOF), from the oral, nontumour and tumour samples of these patients (Table 2). Whole genome short-read sequencing was performed on these isolates, and three were determined contaminated and removed from analyses. Fifty-three recovered genomes belong to *F. nucleatum* (n=38) and *F. periodonticum* (n=15) species complexes, of which phylogenetic reconstruction was performed separately. Core-genome phylogeny of *F. nucleatum* showed that tumour-associated isolates were detected in all four subspecies (*animalis, vincentii, nucleatum, polymorphum*) (Figure 4A). In the *F. periodonticum* phylogeny, tumour-associated isolates (n=8, isolated from P18, P40) formed a distinct cluster that is phylogenetically separated from the available references (Figure 4B). These isolates all showed ∼91% average nucleotide identity (ANI) to the closest *F. periodonticum* references, suggesting that they constitute a novel subspecies of this species complex, denoted herein as novel *F. periodonticum* (novelFperi). Likewise, two gut isolates (H16-13, H16-14) shared 93% ANI to the closest *F. nucleatum* references and were phylogenetically distant from the remaining *F. nucleatum* isolates, potentially indicative of a novel *F. nucleatum* subspecies. Across the two phylogenies, we identified 14 phylogenetic clusters (PCs; 2 – 6 isolates each) and 12 singletons originating from this study’s collection, which were collectively named as PCs herein.

**Table 2.**
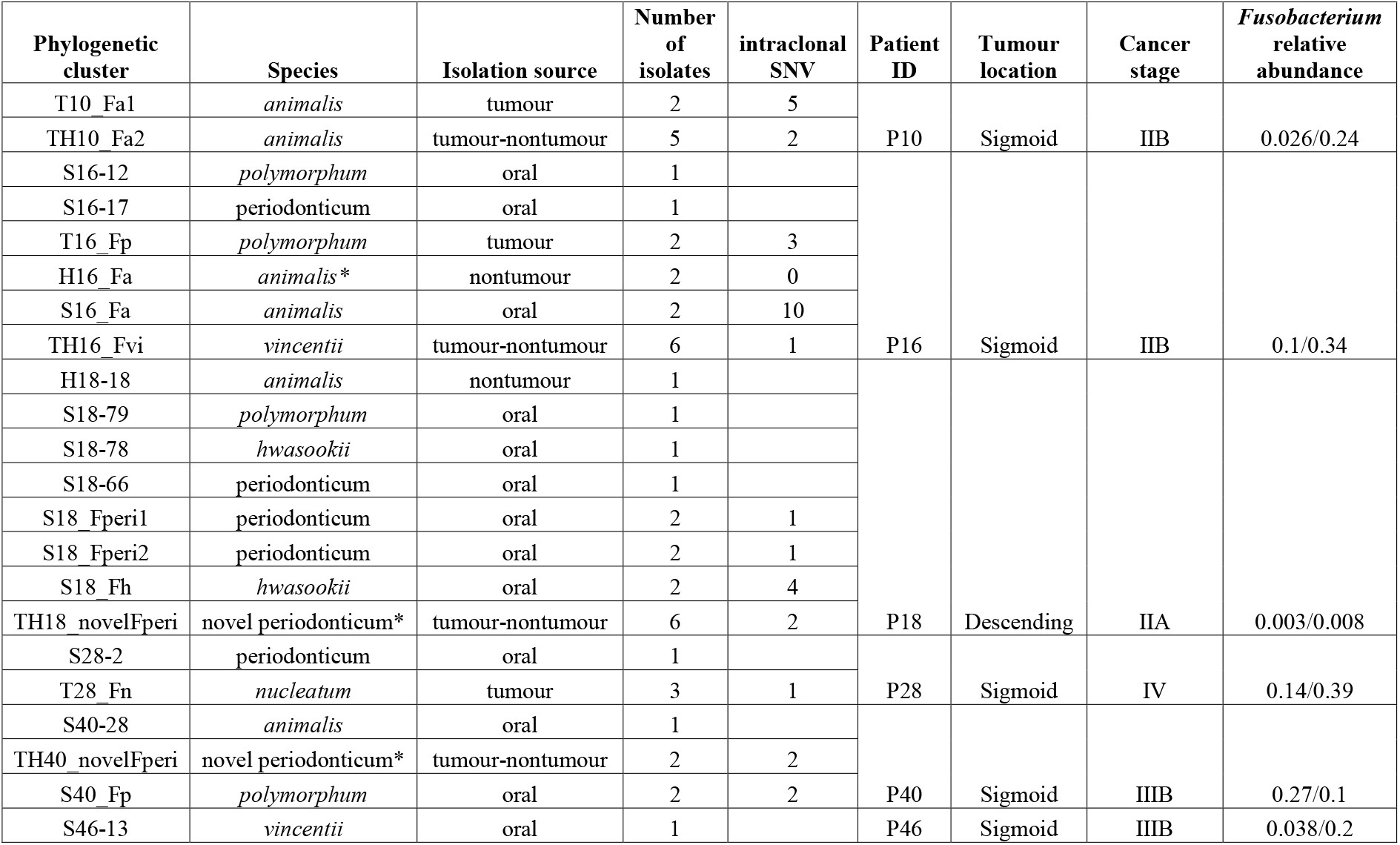
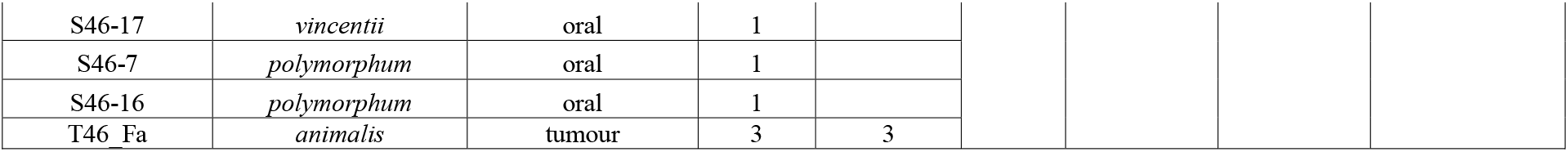
Summary of *Fusobacterium* isolates recovered from this study. Species names in italic represent *Fusobacterium nucleatum* subspecies, and (*) denotes a potential new subspecies. *Fusobacterium* relative abundance, inferred from 16S rRNA gene profiling results, showed the values for non-tumour/tumour samples for each cancer patient. SNV: single nucleotide variant.

**Figure 4.**
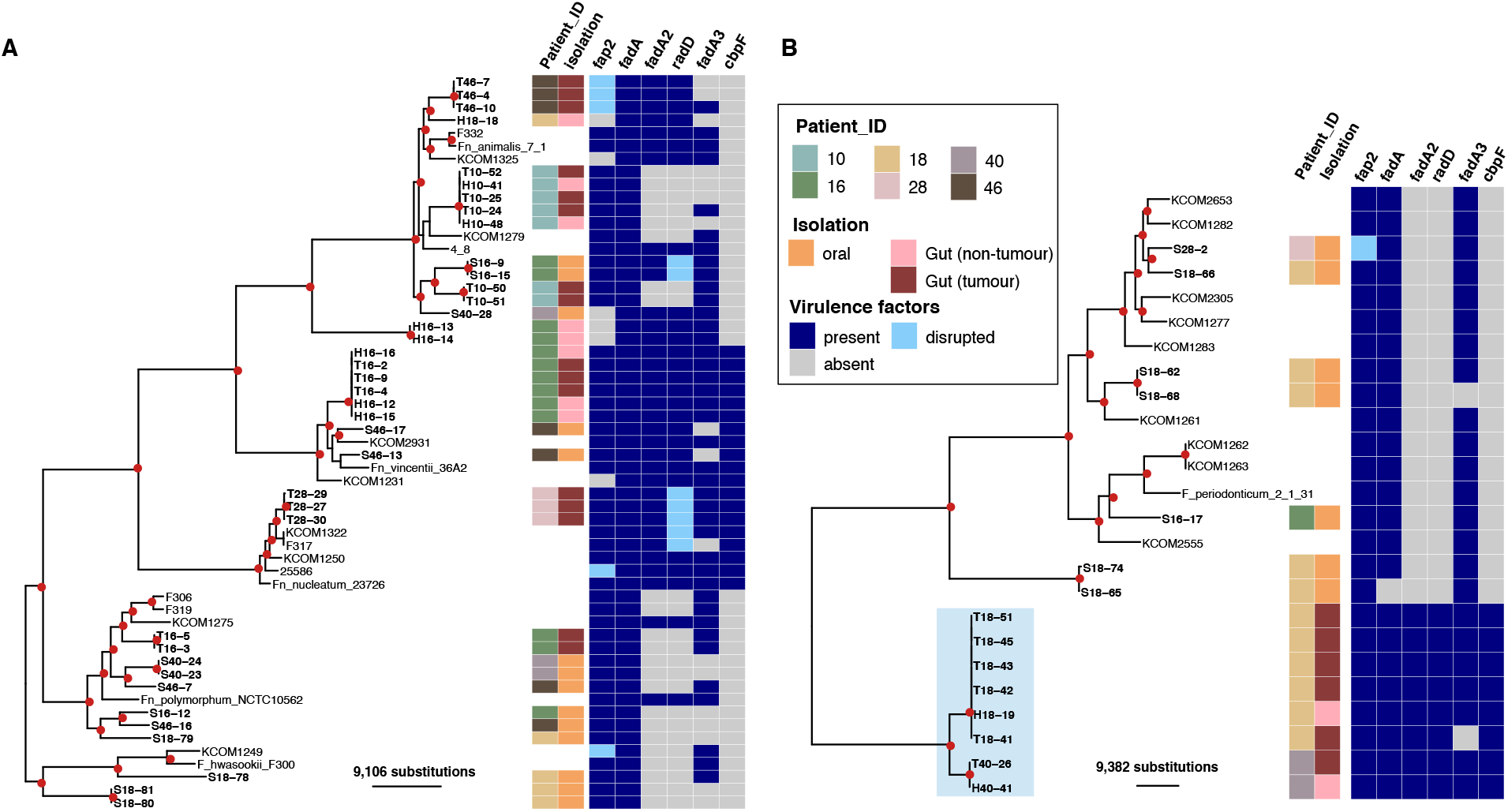
Maximum likelihood phylogenies of *Fusobacterium* isolates from this study. (A) *F. nucleatum* species phylogeny constructed from the alignment of 516 core genes (89,900 variant sites; N=57), using *F. hwasookii* clade as an outgroup. (B) *F. periodonticum* species phylogeny constructed from alignment of 863 core genes (106,738 variant sites, N=26). Red circles at internal nodes denote bootstrap values ≥ 80. The associated metadata on the right describe the patient ID and clinical origin of isolates (where reference genomes were left blank), and the genomic presence of several virulence factors (*fap2, fadA, fadA2, radD, fadA3, cbpF*). Light blue shaded area covers isolates identified as novel *F. periodonticum* subspecies. The scale bars denote the estimated number of substitutions.

The *Fusobacterium* population within each individual patient was diverse (2 – 7 PCs). Several *Fusobacterium* species/subspecies were detected in each patient’s saliva, sometimes with more than one PCs of the same subspecies (P18, P46) (Table 2). Likewise, we observed similar diversity in gut-associated isolates, with more than one PCs detected in three patients (P10, P16, P18). Most patients did not share the same *Fusobacterium* subspecies recovered from both oral- and gut-associated isolates, except for P16 (*polymorphum*). However, phylogenetic evidence confirmed that the two niches harboured distinct populations, which were ∼16,955 SNPs apart (Figure 4A). Particularly, oral *Fusobacterium* isolates from P18 (n=9) belonged to six different PCs (mostly *F. periodonticum* and *F. hwasookii*), while 6/7 gut isolates were of a single novelFperi clone. By contrast, *Fusobacterium* from tumour and nontumour sites were frequently clustered in the same PC (n=4; in P10, P16, P18 and P40), indicating that the same bacterial clones have colonized and spread beyond the tumour microenvironment. We used the mapping approach to confidently inspect the intraclonal variations within these PCs, and showed that they shared minimal genetic differences in the core genome (1 – 2 SNVs). These values fall in range with the variation observed in five other gut PCs (with either tumour or nontumour isolates; 0 – 5 SNVs) and five other oral PCs (1 – 10 SNVs).

### Variation in *Fusobacterium* virulence gene content

We next sought to examine the presence of several *Fusobacterium* virulence factors, of which pathogenicity has been proven in experimental studies, including genes encoding adhesin (*fadA, cbpF*), lectin (*fap2*), and bacterial co-aggregation factor (*radD*) [12,13,29,30]. RadD is an autotransporter facilitating *Fusobacterium’s* interspecies interaction in polymicrobial biofilms [30], while CbpF inhibits CD4^+^ T-cell response through CEACAM1 binding and activation [31]. Genomic screening showed that *fap2* was present and intact in the majority of genomes from both species (49/53), with disruptive mutations occurring in some isolates, such as the tumour-associated *F. nucleatum animalis* in P46 (Figure 4A). We also detected *fadA* in all isolates (except S18-65), with all *F. periodonticum* variants one amino acid shorter (codon A22) than the canonical FadA found in *F. nucleatum* (129 aa). The other elements showed variable presence among the examined genomes. For example, *cbpF* was present in all *F. nucleatum nucleatum, F. nucleatum vincentii*, and novelFperi, while *radD* was co-localised with *fadA2/radA* (a 122 aa *fadA* homolog) in 28 isolates. Another *fadA* homolog (*fadA3*) with unknown function was prevalent in both two *Fusobacterium* species. Phylogenies of FadA and CbpF showed that the two tree topologies were largely in agreement with those inferred from the core genomes, suggesting the absence of horizontal gene transfer (Figure S3). By contrast, the clustering pattern observed in the Fap2 phylogeny was concordant to subspecies classification for *F. nucleatum nucleatum, F. nucleatum vincentii*, and *F. periodonticum*, but was admixed for *F. nucleatum polymorphum, F. hwasookii* and *F. nucleatum animalis* (Figure S4A). *fap2* encodes a very large protein of variable length (median of 3938 aa [range: 3436 – 4669]), and the protein length showed some correlation with its phylogenetic clustering, with variants > 4200 aa (n=6) all belonging to a monophyly composed of *F. hwasookii* and *F. nucleatum polymorphum*. Similarly, the RadD phylogeny did not concur with those inferred from the core genomes, and its length variation (median 3526 aa [range: 3461 – 3602]) also showed association with the tree topology (Figure S4B). *radD* was ∼800 bp downstream of *fadA2*, which is flanked by an IS150 transposase on the *F. nucleatum* 23726 reference genome. This could explain the mobilization mechanism of *radD*-*fadA2* across the *Fusobacterium* phylogeny. These data indicate that the autotransporter encoding genes *fap2* and *radD* may have undergone frequent horizontal gene transfer or recombination in the *F. nucleatum* species complex.

## Discussion

Our study revealed the composition of microbiome perturbations at the tumour mucosa of Vietnamese patients with CRC and non-cancerous colorectal polyps. Tumour-enriched taxa include mostly bacteria of putative oral origin, such as *F. nucleatum, Leptotrichia, Gemella, C. rectus*, and *Selenomonas*, which agrees with findings from previous studies profiling either gut mucosal or faecal microbiomes in different CRC populations [8,9,32]. This suggests that the proliferation of oral bacteria at the gut mucosa could be a universal signature of CRC microbiomes. Several of these oral taxa shared identical ASVs between the oral and gut niches, pointing to the oral origin of tumour-associated taxa. Our analysis found that these bacteria also display a co-occurrence pattern, indicating that they likely co-exist in a biofilm-like aggregate upon colonization at the gut mucosa. Indeed, previous research has confirmed the frequent presence of polymicrobial biofilms composed of oral taxa (*F. nucleatum, Peptostreptococcus, Gemella*) in colorectal tumour tissues [33]. Among the oral bacteria, *F. nucleatum* stands out for its ability to form “bridging” interactions with other bacteria via the presence of several adhesins [11]. *F. nucleatum* was recently reported to secrete FadA with amyloid properties, which confers acid tolerance and provides a scaffold for biofilm formation [34]. In addition, our analyses pointed to the significant presence of *Leptotrichia* in tumour mucosa, especially in advanced tumours. This association, however, has only been noted in few studies [32,35]. This may be due to the differences in sampling location, as tumours excised from the distal colon (as performed for all cases in our study) were reported to harbour a higher abundance of *Leptotrichia*, compared to those originating from the proximal colon [35]. The overabundance of *Leptotrichia* in the salivary microbiome has been implicated in patients with malignant oral leukoplakia and pancreatic cancer [36,37], as well as with CRC as shown in this study. *Leptotrichia* belongs to the same order as *Fusobacterium* (Fusobacteriales) and could carry virulence factors similar to those found in the latter genus. It is noteworthy that the predominant tumour-associated *Leptotrichia* taxon (ASV-13) could be detected from different CRC patients, but was in very low abundance in these patients’ salivary microbiomes. This suggests that a distinct *Leptotrichia* species/genotype was associated with CRC, which warrants more in-depth investigations.

Asides from oral taxa, *Hungatella* overabundance was the most significant signature of CRC microbiome in our dataset. This falls in line with results from a recent metagenomic meta-analysis, showing that *Hungatella hathewayi*’s specific choline trimethylamine-lyase gene (*cutC*) was significantly enriched in the faecal microbiomes of CRC patients [10]. Moreover, colonic *H. hathewayi* could induce hypermethylation in prominent tumour suppressor genes, thus silencing their functions and promoting intestinal epithelial cell proliferation [38]. Combining quantitative detection of microbial markers (*H. hathewayi, F. nucleatum, Lachnoclostridium*, and *Bacteroides clarus*) with faecal immunochemical test greatly increased sensitivity (reaching ∼94%) for diagnosing CRC in a Chinese population [39]. Consistent with this finding, our analyses suggested that *Hungatella* and *Lachnoclostridium* were overabundant and co-occurring in the tumour mucosa of the Vietnamese cohort, which supports the feasibility of applying such microbial detection test for non-invasive CRC screening in our setting. On the other hand, we found that *F. mortiferum* was the most significantly enriched taxon in the polyp control group. *F. mortiferum* was known as a hallmark for dysbiosis in infectious diarrhoea [40], and recent studies have also reported the abundance of *F. mortiferum* in patients with colorectal polyps [41,42]. Furthermore, this species was shown to be present in the gut microbiomes of ∼60% of a cohort in Southern China, albeit in very low abundance (∼0.5%) [43]. Unlike other *Fusobacterium* species, *F. mortiferum* was devoid of distinctive virulence factors such as adhesins FadA and Fap2 [44], but could utilize a wide range of sugars for growth independent of amino acid metabolism [45]. The association between *F. mortiferum* and colorectal polyps will need to be further addressed in future studies.

Despite the increasing importance of *F. nucleatum* in the pathogenesis of CRC and other invasive diseases [11], genomic characterization of these bacteria from patient populations is currently limited due to technical difficulties in *Fusobacterium* isolation. Here, we applied targeted culturomics approach, which combines anaerobic culturing, high-throughput identification by MALDI-TOF and WGS, to study the *Fusobacterium* population in high resolution and help uncover novel bacteria [46]. Indeed, we discovered novel subspecies of both *F. nucleatum* and *F. periodonticum* from culturing the gut mucosa, showing that the microbiomes in non-Western settings offer untapped diversity. Using metagenomic assemblies from Chinese faecal microbiomes, Yeoh and colleagues have proposed several new *Fusobacterium* species (based on 95% ANI cutoff) [44]. However, our WGS approach provided more accurate and complete realization of the bacterial genomes, which contributes to the global representation of *Fusobacterium* diversity (with 26 non-duplicate assemblies added). Furthermore, our approach allows for delineation of bacteria from tumour and non-tumour sites, which is inaccessible by faecal metagenomes. The populations of *Fusobacterium* colonizing the oral cavity and gut mucosa were heterogeneous within each individual, even at the subspecies level, which mirrors the diversity observed previously for gut commensals such as *Bifidobacterium* species [47]. Though this study did not provide evidence of genetic relatedness between oral and gut *Fusobacterium* isolates, this likely point to the high diversity of *Fusobacterium* in the oral niche [48]. Besides, *Fusobacterium* is abundant in subgingival dental biofilms, which our salivary sampling did not fully capture. Previous research deploying WGS has demonstrated that oral and tumour-originated *F. nucleatum* shared little genetic divergence, supporting the notion that oncogenic *Fusobacterium* arise from the patient’s oral microbiome [49]. Chronic infections with *Helicobacter pylori* at the stomach, which increases the risk of gastric cancer, usually result in extensive clonal propagations detected by WGS within each patient, though isolates were collected in a single timepoint [50]. This prolonged colonization scenario contrasts with our observations in CRC, in which multiple *Fusobacterium* clones (with minimal intraclonal variation) were present at each patient’s gut mucosa. Given that CRC could take years to develop, we speculate that the *Fusobacterium* population at the tumour site fluctuates in response to the frequent seedings from the highly diverse oral source. Longitudinal study design is needed to address this hypothesis, and to further assess how *Fusobacterium* adapts to the gastrointestinal pressure.

The two well-described major virulence genes (*fadA* and *fap2*) were identified in the majority of *Fusobacterium* genomes, regardless of niche. This concurs with previous research reporting the high prevalence of *fadA* and *fap2* in *F. nucleatum* and *F. periodonticum* metagenomic assemblies from a cohort in China [44]. These suggest that *Fusobacterium* with high virulence potential are prevalent in the human population, and the genetic presence of *fadA* and *fap2* is not suitable for predicting the risk of *Fusobacterium*-related CRC. All gut-derived novelFperi isolates harboured the examined virulence genes (*fadA, fap2, radD*, and *cbpF*), which was more similar to *F. nucleatum* compared to *F. periodionticum*. Moreover, *fap2* and *radD* showed variation in gene length and evidence of horizontal gene transfer, underlying the significance of dynamic evolutionary processes in shaping *Fusobacterium*’s virulence landscape. Since Fap2 orchestrates *F. nucleatum* invasion into CRC tumour cells via specific binding to Gal-GalNAc, this ligand-receptor interaction was recently proposed as a target for clinical intervention in *Fusobacterium*-enriched CRC [51]. Interestingly, our genetic analysis predicted that *fap2* was either missing or truncated in some gut-associated *Fusobacterium* isolates, which may indicate the complex lifestyle of *Fusobacterium* once colonizing the gut environment.

Some limitations were notable in our study design. Due to ethical concerns, patients with colorectal polyps were selected as the control group, instead of healthy age-matched individuals. Our interpretations do not extend to cancer in the proximal colon, though previous reports have noted that proximal CRC tumours had a higher *Fusobacterium* abundance [52]. The sample size of cultured *Fusobacterium* isolates was moderate and did not include longitudinal sampling, so it was not possible to investigate the bacterial evolution in longer timeframe. Notwithstanding these shortcomings, our study reconfirmed the prominent role of oral anaerobic conglomerates in CRC microbiome in an understudied Asian population, and provided new insights into the genomic diversity of the oncobacterium *Fusobacterium*. The observed diversity in this organism should be taken into account when designing future diagnostic or therapeutic tools that target *Fusobacterium*.

## Material and Methods

### Study design and sample collection

This prospective case-control study enrolled adult Vietnamese patients (≥18 years old) admitted at Binh Dan Hospital, a large surgical hospital in Ho Chi Minh City Vietnam, from December 2018 to January 2020. The study received ethical approval from the Ethics Committee of Binh Dan Hospital. Written informed consent was obtained from all study participants. Cases were defined as patients diagnosed with left-sided colorectal cancer (distal colon and rectum) of stage II onward, who received colectomy treatment and underwent non-antibiotic pre-operative bowel preparation. Controls were patients diagnosed with colorectal polyps (single/scattered non-cancerous polyps at distal colon or rectum), who received polypectomy at the hospital. For both cases and controls, patients were excluded if they (1) had received antimicrobial treatments within two weeks prior to enrolment, (2) had additional gastrointestinal infections or obstructions, or (3) were immunocompromised. Additionally, the study excluded patients who had received chemo- and/or radio-therapy within four weeks prior to enrolment (for cases) and those were diagnosed with familial adenomatous polyposis (controls).

Demographic and clinical information were collected from study participants at recruitment. The calculated body mass index (BMI) was categorized based on WHO recommendation for Asian populations [53]. Cancer stage classification was based on the TNM Staging system [54]. A saliva sample (∼3mL) was collected pre-operation from each study participant (by spitting into a sterile container). For cases, the mucosa epithelia at the tumour and adjacent non-tumour (2-10 cm away from the tumour) sites were collected aseptically from the excised colon. For controls, we collected colorectal polyps and 2-3 biopsies of non-polyp mucosal epithelium (∼50 mg) during colonoscopy. All clinical samples were stored on ice and transported back to the OUCRU laboratory within 4 hours, then were stored in -80°C until further experiments.

### 16S rRNA gene sequencing

We selected 43 cases and 25 controls for microbiome profiling. Total DNA was extracted from the gut mucosa samples (n=136) using the FastDNA spin kit for soil (MP Biomedicals, USA), with bead-beating step on Precellys 24 homogenizer (Bertin Instruments, France). DNA from the saliva samples (n=67, one missing) was extracted using the ReliaPrep Blood gDNA Miniprep (Promega, USA). For microbiome profiling, all samples underwent primary PCR amplification (30 cycles) using the conventional V4 primers (515F-806R) and KAPA HiFi Hot Start DNA polymerase (KAPA Biosystems, USA), and secondary PCR was performed to add dual-indexes (IDT, USA) to each sample, following procedures optimized in a published protocol [55,56]. Additionally, we applied the same procedures to a positive control (eight species Zymo mock community, Zymo Research, USA) and six negative controls (two for each DNA extraction kits, and two no-template PCR amplifications), in order to respectively evaluate the experiment’s efficacy and detect contamination from reagents and kits (kitome) [57]. 16S rRNA sequencing was performed for all samples on one run of the Illumina MiSeq platform, to generate 250 bp paired-end reads.

### Microbiome data analysis

All data analyses were conducted in R (v4.1.1) and Rstudio using multiple packages, including ‘dada2’, ‘phyloseq’, ‘DESeq2’, ‘ANCOMBC’, ‘corncob’, ‘philr’, ‘ggplot2’, ‘vegan’, ‘SpiecEasi’ and others [24– 26,28,58–61]. Generated sequence reads were analysed under the amplicon sequence variant framework (ASV) using DADA2, in which statistically denoised forward and reverse read pairs were merged to create error-corrected ASVs with single-nucleotide resolution [62,63]. We retained ASVs with length ranging from 249 to 256 bp, which matches the desired length of the amplified V4 region, and chimeric sequences were detected and removed independently for each sample. Taxonomic assignment (up to the species level) was performed using the RDP Naïve Bayesian Classifier implemented in ‘dada2’ package, on the SILVA v138 train dataset [64]. Further filtering removed ASVs matching the following criteria (1) classified as ‘Mitochondria’ or ‘Archaea’, (2) unclassified at Kingdom or Phylum level, (3) identified as kitome or contamination from mock community (except *Escherichia* and *Enterococcus* ASVs), or (4) identified as low abundant singletons (abundance ≤10 counts and present in only one sample). This resulted in 2,461 ASVs detected across 203 samples (68 participants), totalling 5,250,754 sequences.

Saliva and gut mucosal microbiomes were then analysed separately. For saliva microbiomes, we removed singleton ASVs with abundances < 79 sequences and one sample with low sequencing depth (837 sequences). The filtered ASVs (n=865) were aligned using PASTA [65], and a maximum likelihood phylogeny was constructed under the GTR+G model using IQ-Tree (with 1,000 rapid bootstrap) [66]. The resulting phylogeny was used to transform the ASV count matrix into isometric log-ratio (ILR) ‘balances’ (weighted log-ratio between two ASVs), using the “philr” package (with zero counts imputed using the “cmultRepl” function) [60,67]. This transformation allowed statistical analyses to be performed accurately in the Euclidean space [68]. Ordination was performed using principle coordinate analysis (PCoA) on a calculated Euclidean distance matrix, implemented in package ‘phyloseq’. To identify covariates which explain the salivary microbiome structures, we performed redundancy analysis on the ‘balance’ value matrix of 62 samples with complete metadata (case-control grouping, age, sex, BMI, presence of oral diseases, and status of active smoking in the last two years). We repeat the same analytical procedures on the gut mucosal microbiome data. Low-abundance singleton ASVs (< 44 sequences) and seven samples with low sequencing depth (<1,300 sequences each) were removed, retaining 1,073 ASVs across 129 samples for downstream analyses. We tested the association between covariates and the gut mucosal microbiome structures using redundancy analysis, performed on the ILR-transformed ‘balance’ values of 120 samples with complete metadata (sample type, age, sex, BMI, history of diarrhea in seven days, diabetes, high blood pressure, active consumption of alcohol in the last two years, and anatomical location of samples). The ILR-transformed values were used to calculate the differences among the samples (beta-diversity), within and between participants. In addition, the gut mucosal microbiomes (n=129) were clustered into community state types (CSTs) using the partition around medoid (pam) algorithm on the calculated ILR-transformed distance matrix, with the optimal number of CSTs (k=2) determined by gap statistic and average silhouette width (asw) [69]. The random forest classification algorithm (10,000 trees) was then used to evaluate this clustering performance (in term of error rate) and to identify ‘balances’ differentiating the two CSTs, using the ‘rfsrc’ function implemented in package ‘randomforestSRC’ [70].

### Evaluating differential abundances

In order to detect ASVs that showed significantly differential abundance between two examined groups, we utilized the compositional data analysis approach implemented in ANCOMBC [24]. In addition, the same comparisons were performed using DESeq2 and corncob to check for consistent results, as recommended in recent benchmark studies [71,72]. The comparisons include salivary microbiomes in cases (n=43) and controls (n=23); paired tumours (n=43) against adjacent non-tumours (n=43); paired polyps (n=16) against non-polyp biopsies (n=16); tumours (n=43) against non-polyp biopsies (n=24); tumours of cancer stage III-IV (n=24) against stage II (n=18). For paired comparison within cases and controls, the model design was set to “∼Patient + sample_type” to increase statistical power [73]. Multiple hypothesis testing was corrected using Holm or Benjamini-Hochberg method, setting false discovery rate as 0.05. ANCOMBC and corncob approaches were carried out using default parameters. For DESeq2, count data were normalized using either the native negative binomial distribution (saliva, tumours vs. biopsies) or the zero-inflated negative binomial (ZINB) distribution implemented in the package ‘zinbwave’ (paired samples in cases and controls, between cancer stages) [74]. Library size corrections were estimated using DESeq2’s ‘poscounts’ method. All comparisons were performed using likelihood ratio test, and ASVs with adjusted p-value < 0.05 (and base mean >20 for DESeq2) were considered significant hits. To minimize the number of false positives, ASVs which showed significant hits in at least two tested methods were considered differentially abundant and included in final interpretation. We manually compared the ASV sequences of interest to the expanded Human Oral Microbiome Database (HOMD; www.homd.org/), and species identification was assigned if the ASV showed >99% nucleotide similarity to that in the database.

### Correlation network

We constructed a correlation network of gut mucosal microbiomes from colorectal cancer patients (n=86), using 117 most representative ASVs, defined as ones with abundance of at least 10 sequences detected in at least 15 samples. This filtering resulted in a median sample retainment rate of 77% [70% -85%]. Zero counts were imputed using the “CmultRepl” function, and the correlation network was constructed using CCLasso, with 250 bootstrap and three-fold cross validation [27]. Interactions with adjusted p values < 0.01 and absolute correlation strength > 0.37 were considered significant hits. Additionally, a separate correlation network was inferred using SpiecEasi (Meinshausen-Buhlmann’s neighbourhood selection, nlambda=20) on the same dataset [28]. Both these methods have been demonstrated to produce robust performance in a recent benchmark study [75]. To avoid spurious hits, only significant interactions detected by both the CCLasso and SpiecEasi approaches were included in the final visualization. We applied the same procedures to construct correlation networks of microbiomes in saliva samples (n=66, 115 ASVs) and controls’ gut mucosa (n=43, 90 ASVs).

### Fusobacterium isolation and whole genome sequencing

*Fusobacterium* isolation was performed on six selected case patients (P10, P16, P18, P28, P40, P46), whose *Fusobacterium* relative abundance in the tumour microbiome exceeded 0.5% as inferred by microbiome profiling. The respective samples (saliva, tumour, and non-tumour mucosa) were subjected to anaerobic culturing in a Whitley A35 anaerobic workstation (Don Whitley Scientific, UK) supplied with 5% CO_2_, 10% H_2_, and 85% nitrogen gas, following the *Fusobacterium* isolation procedures established previously [76]. Briefly, mucosal tissues were thawed on ice, and ∼100 mg tissues were aseptically excised and anaerobically homogenized in Diluent A to maximize bacterial recovery. The suspension (100 µL) was then plated onto the selective media at two-fold and four-fold dilutions (EG agar supplemented with L-cysteine HCl, 50 ml/liter of defibrinated sheep blood, 7 mg/liter of crystal violet, 5 mg/liter of vancomycin, 30 mg/liter of neomycin, and 25 mg/liter of nalidixic acid; Sigma-Aldrich, Germany). Thawed saliva samples were plated directly on the selective media, either undiluted or at two-fold dilutions. Plates were incubated at 37°C for 48-72h, and colonies (up to 10) resembling that of *Fusobacterium* were picked from each plate and sub-cultured on new EG media to confirm purity and select for single colonies. The isolate’s taxonomic identities were queried using MALDI-TOF, and those characterized as *Fusobacterium* species were retained. A total of 56 *Fusobacterium* isolates were recovered and subjected to DNA extraction using the Wizard genomic extraction kit (Promega, USA). For each isolate, 1 ng DNA was used to prepare the sequencing library using the Nextera XT library preparation kit, following the manufacturer’s instruction. Normalized libraries were pooled and sequenced on an Illumina MiSeq platform to generate 250 bp paired-end reads.

### Pangenome analysis, phylogenetic reconstruction and screening for virulence genes

FASTQC was used to check the sequencing quality of each read pair [77], and Trimmomatic v0.36 was used to trim sequencing adapters and low-quality reads (paired end option, SLIDINGWINDOW:10:22, HEADCROP:15, MINLEN:50) [78]. For each isolate, the trimmed read set was input into Unicycler v0.4.9 to construct the de novo assembly, using default parameters, and contigs of size over 500bp were retained for downstream analyses [79]. The assemblies were checked for traces of contamination using Checkm, and three assemblies were shown contaminated and discarded [80]. The resulting assemblies were of adequate quality, with median size of 2,125,169 bp [IQR: 2,067,843 – 2,168,429], median number of contigs of 133 [IQR: 86 – 173] and the median N50 of 35,535 bp [IQR: 21,685 – 51,953]. Prokka v1.13 was used to annotate the assemblies, using the well-annotated *F. nucleatum* 23726 (accessed via FusoPortal) as reference [81]. To provide preliminary taxonomic classification up to the subspecies level, FastANI was used to calculate the average nucleotide identity (ANI) between the individual assembly and a set of *Fusobacterium* references, with an ANI value ≥ 95% denoting a shared species/subspecies [82]. The pangenomes of 57 *F. nucleatum/hwasookii* isolates (38 sequenced herein plus 19 references) and 25 *F. periodonticum* isolates (15 sequenced herein plus 10 references) were constructed separately using panX, which clusters orthologous proteins based on individual gene tree construction and adaptive postprocessing instead of relying on fixed nucleotide identity cutoff [83]. The respective core genome from each species complex was aligned, with invariant sites removed, producing SNP alignments of 89,900 bp (*F. nucleatum/hwasookii* complex) and 106,738 bp (*F. periodonticum* complex). These were input into RAxML to construct maximum likelihood phylogenies, under the GTRGAMMA substitution model with 300 rapid bootstraps [84]. Using the pangenome analysis output (clustered orthologous proteins and gene presence/absence matrix), we screened for the presence of several known *Fusobacterium* virulence genes (*fap2, fadA, radD, cbpF*). The intact presence or synteny of each genetic element was checked manually by gene alignment (Seaview) or genome visualization (Artemis) tools [85,86]. Since Fap2 is a large protein (3,000 – 4,600 amino acid) with varying size among the *Fusobacterium* species/subspecies, we checked for its intactness using ARIBA and mapping approach, with clone-specific *fap2* serving as mapping reference [87]. Visualization of phylogenetic tree and associated metadata was performed using package ‘ggtree’ [88]. Individual protein sets were aligned and inspected in Seaview, and phylogenies were constructed in RAxML, using the PROTGAMMAGTR model and 200 rapid bootstraps.

### Intra-clonal variation examination

To investigate intra-clonal variation with high confidence, we examined single nucleotide variants (SNV) among isolates belonging to the same phylogenetic cluster (Figure 4 and Table 2), using the mapping approach recommended previously [89]. For each phylogenetic cluster, trimmed fastq files from the isolates were concatenated and input into Unicycler to construct a pan-assembly, with contigs less than 500 bp removed. This pan-assembly was ordered against an appropriate *Fusobacterium* reference using ABACAS, depending on its respective species/subspecies, creating a pseudogenome reference [90]. Trimmed paired-end reads from each isolate were mapped against this reference using a custom wrapper script. Briefly, mapping was conducted using BWA MEM algorithm and samtools v1.8 [91,92], with duplicate reads removed using PICARD, followed by indel realignment by GATK [93]. Reads with nonoptimal local alignment were subsequently removed using samclip (--max 50; github.com/tseemann/samclip), to avoid false positives during variant calling. SNVs were detected using the haplotype-based caller Freebayes [94], and low quality SNVs were removed using bcftools if they met any of the following criteria: consensus quality < 30, mapping quality < 30, read depth < 4, ratio of SNVs to reads at a position (AO/DP) < 85%, coverage on the forward or reverse strand < 1 [95]. Mapping coverage at each position was inferred using samtools “depth” command (-a -Q 30). The bcftools ‘consensus’ command was used to generate a pseudosequence (with length identical to the mapping reference), integrating the filtered SNVs and invariant sites, and masking the low mapping region (depth < 4) and low-quality SNVs with ‘N’. The presence of high quality SNVs were validated by manual visualization of output bam files in Artemis, and SNV pertaining to recombination, transposons, plasmids, or repetitive elements were excluded from interpretation.

## Data availability

Raw sequence data are available in the NCBI Sequence Read Archive, including ones for 16S rRNA sequencing (BioProject PRJNA791834) and *Fusobacterium* whole genome sequencing (BioProject PRJNA791829). Source data and R codes used for the microbiome analysis and visualization are deposited in Github (https://github.com/Hao-Chung/Vietnam_CRC_microbiome).

## Funding information

HCT is a Wellcome International Training Fellow (218726/Z/19/Z). LJH is supported by Wellcome Trust Investigator Awards 100974/C/13/Z and 220876/Z/20/Z; the Biotechnology and Biological Sciences Research Council (BBSRC), Institute Strategic Programme Gut Microbes and Health BB/R012490/1, and its constituent projects BBS/E/F/000PR10353 and BBS/E/F/000PR10356. SB is a Wellcome Senior Research Fellow (215515/Z/19/Z).

## Disclosure of potential conflicts of interest

The authors report no potential conflicts of interest.

## Consent for publication

This study does not publish any identifiable details related to the participant.

## Acknowledgements

The authors wish to thank all participants and their caretakers for their participation in the study, and the pathology laboratory of Binh Dan Hospital for the tumour/biopsy pathology results.

## Notes

### Competing Interest Statement

The authors have declared no competing interest.

