## Supplementary figures for "Mucosal microbiomes and *Fusobacterium* genomics in Vietnamese colorectal cancer patients"

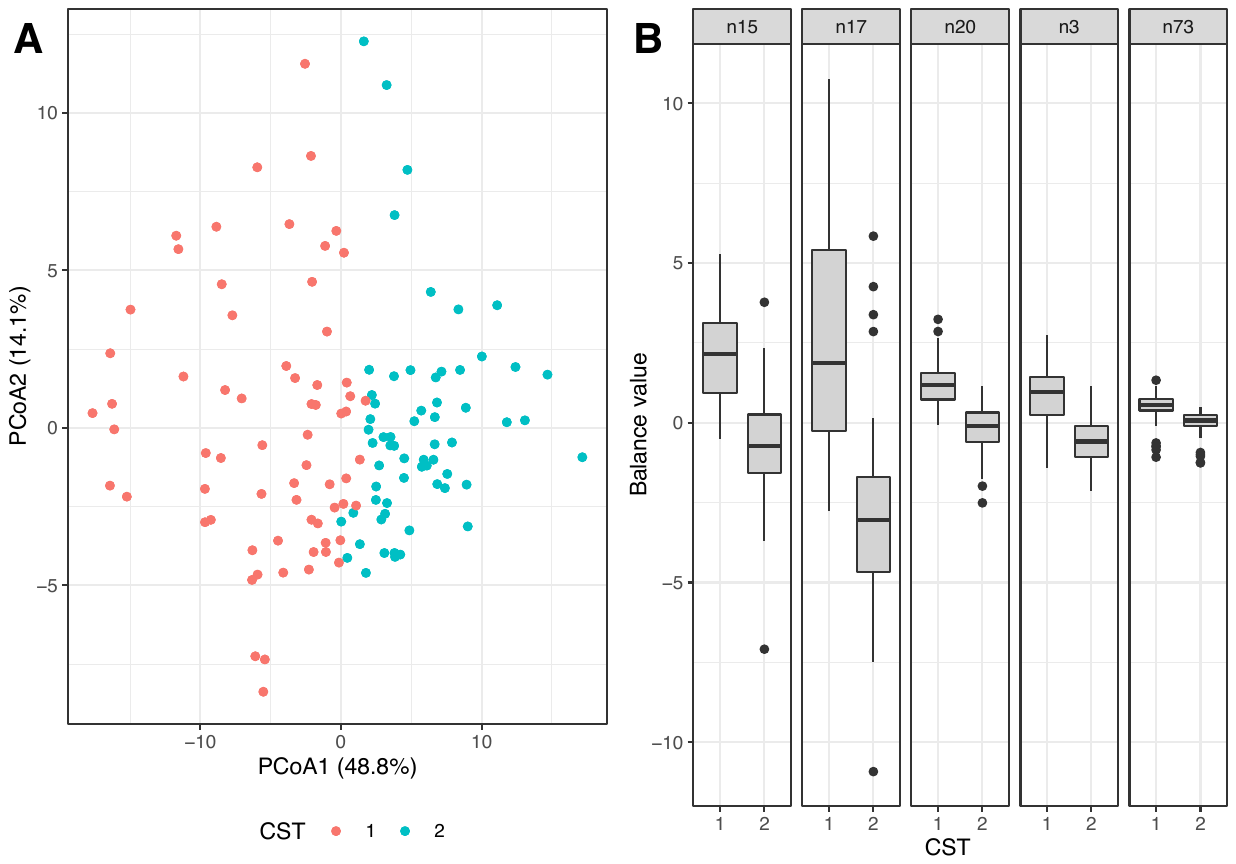

**Figure S1** Clustering of the gut mucosal microbiomes into two community state types (CSTs). (A) Principal coordinate analysis (PCoA) conducted on phylogenetic-assisted isometric log-ratio (PhILR) transformed data (similar to Figure 1B), with samples coloured by CST membership. (2) “Balances” identified by Random forest classification as most important in differentiating the two CSTs, with values shown in boxplot for each CST. N15: Actinobacteria+Proteobacteria/Firmicutes+Fusobacteria, n17: Proteobacteria/Actinobacteria, n20: Gammaproteobacteria/Campilobacterota+Desulfobacterota; n3: other bacteria/Lachnospiraceae, n73: *Escherichia*/*Morganella*.

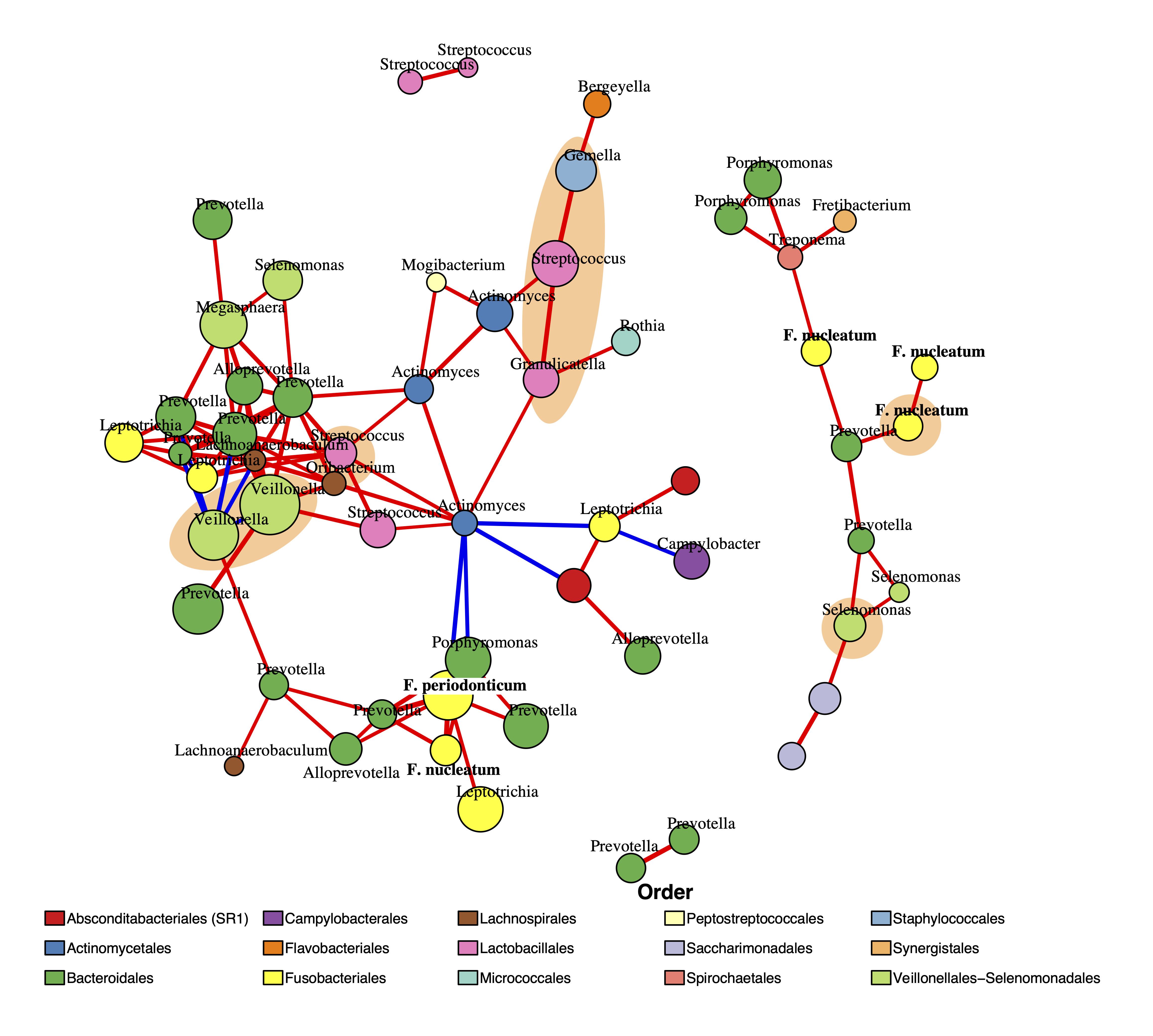

**Figure S2** Correlation network of salivary microbiomes from all participants. The network was constructed from 115 most representative ASVs sampled from 66 mucosal microbiomes, outlining significant interactions detected by both CCLasso (p value ≤ 0.01 and absolute correlation strength > 0.4) and SpiecEasi. Positive and negative interactions were coloured as red and blue lines respectively, with line weight proportional to correlation strength. The ASVs (nodes) were coloured based on taxonomic family (see Legend), with sizes proportional to their relative abundances. The orange shaded area entails ASVs detected in the gut mucosal microbiome correlation network of CRC patients (see Figure 3).

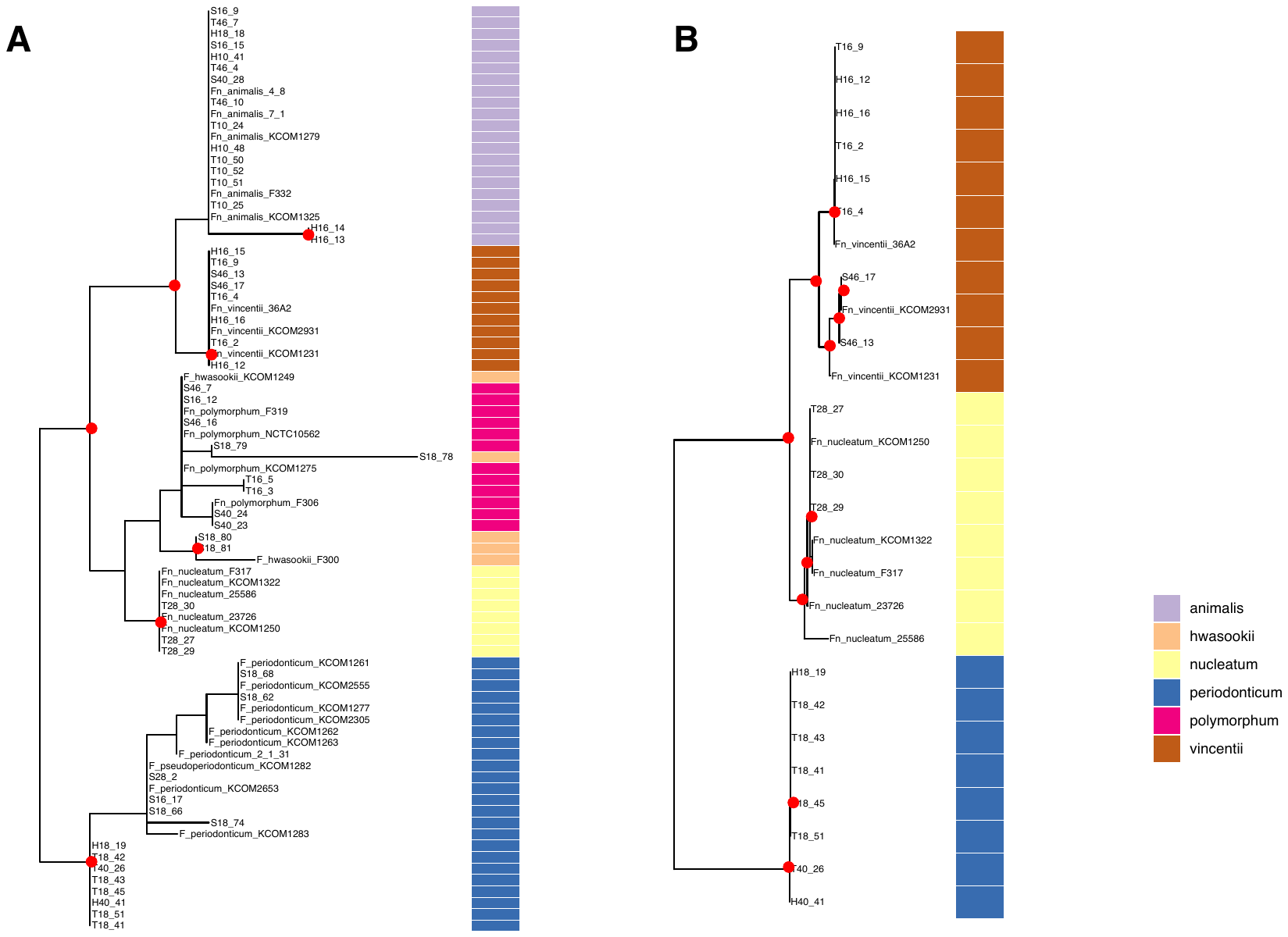

**Figure S3** Maximum likelihood phylogenies of (A) FadA and (B) CbpF proteins. Red circles at internal nodes denote bootstrap values ≥ 70. The column on the right denotes different *Fusobacterium* species/subspecies.

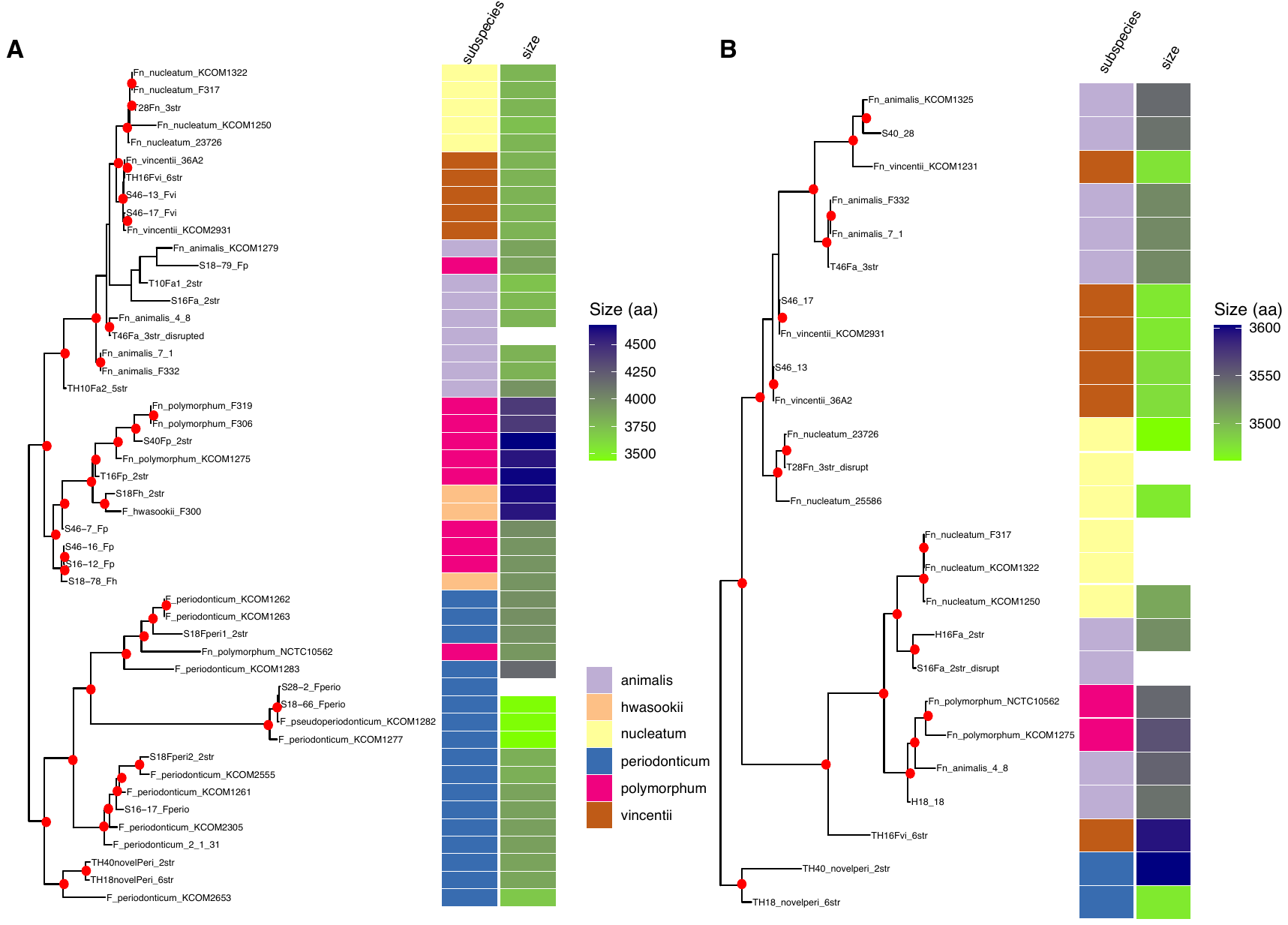

**Figure S4** Maximum likelihood phylogenies of (A) Fap2 and (B) RadD proteins. Red circles at internal nodes denote bootstrap values ≥ 70. The columns on the right denote different *Fusobacterium* species/subspecies and the size of each protein. Disrupted proteins were left blank in size.
